# Altered structural connectivity and functional brain dynamics in individuals with heavy alcohol use

**DOI:** 10.1101/2023.11.27.568762

**Authors:** S. Parker Singleton, Puneet Velidi, Louisa Schilling, Andrea I. Luppi, Keith Jamison, Linden Parkes, Amy Kuceyeski

## Abstract

Heavy alcohol use and its associated conditions, such as alcohol use disorder (AUD), impact millions of individuals worldwide. While our understanding of the neurobiological correlates of AUD has evolved substantially, we still lack models incorporating whole-brain neuroanatomical, functional, and pharmacological information under one framework. Here, we utilize diffusion and functional magnetic resonance imaging to investigate alterations to brain dynamics in *N* = 130 individuals with a high amount of current alcohol use. We compared these alcohol using individuals to *N* = 308 individuals with minimal use of any substances. We find that individuals with heavy alcohol use had less dynamic and complex brain activity, and through leveraging network control theory, had increased control energy to complete transitions between activation states. Further, using separately acquired positron emission tomography (PET) data, we deploy an *in silico* evaluation demonstrating that decreased D2 receptor levels, as found previously in individuals with AUD, may relate to our observed findings. This work demonstrates that whole-brain, multimodal imaging information can be combined under a network control framework to identify and evaluate neurobiological correlates and mechanisms of AUD.

## 1 Introduction

Alcohol use disorder (AUD) is a long-term and recurring neurological condition that can continue unabated despite significant adverse effects on the person, their family, and the broader community. However, the root neurobiological causes of AUD remain undefined, there are limited effective treatment methods available, and relapse rates are around 60% [1]. Significantly, it’s been observed that only a fraction of individuals who regularly consume addictive substances eventually develop a substance use disorder (SUD). This emphasizes the urgent need to uncover biological elements that predispose a person to develop SUDs, and to improve prevention and treatment paradigms.

Individuals with an SUD may be vulnerable because of genetics, developmental differences, hormones, life experiences, environmental and/or adverse social exposures [2]. The brain’s reward circuitry, stimulated by most addictive drugs, depends greatly on dopamine signaling, particularly in the ventral tegmental area (VTA) and dorsal striatum, including the nucleus accumbens (NAc). Chronic exposure to dopamine-stimulating drugs, such as alcohol, can trigger glutamatergic-mediated changes in the striato-thalamo-cortical (specifically orbitofrontal and anterior cingulate cortex - ACC) and limbic pathways (amygdala and hippocampus) that in certain individuals can lead to transition from goal directed to habitual control over drug-seeking behaviors [3]. Several positron emission tomography (PET) studies have revealed that people with SUD of alcohol [4], cocaine [5], heroin [6] and methamphetamine [7] have reduced concentrations of dopamine receptors. One hypothesis is that individuals with lower dopamine receptor levels, due to genetics and/or because of their environment or life experiences, have less than usual dopamine-mediated pleasure from everyday life and therefore may be susceptible to habitual seeking of drug-induced increases in dopamine.

Neuroimaging studies have begun to reveal differences in brain structure and function in individuals with SUDs. A recent meta-analysis revealed brain structures involved across levels of use (SUD vs occasional vs long-term) and substance type, including the thalamus, insula, inferior frontal gyrus, and superior temporal gyrus [8]. Further neuroimaging evidence points to possible reduction in top-down inhibitory control of bottom-up signaling [9], which may support the proposed hypothesis of SUD as a disease of control dynamics [10]. In susceptible individuals, certain stimuli (bottom-up signals) may activate strong urges that in others would be suppressed by top-down inhibition, but in susceptible individuals result in compulsive behavior [11]. Together, the current evidence points toward neurobiological mechanisms of SUDs, which likely involve differences in receptor concentration/function, brain activity patterns and anatomy (gray and white matter [12]). However, a unifying computational model integrating multi-modal observations into a single framework has not been proposed, no doubt hampering our ability to understand the neurobiological mechanisms of SUDs, which in turn is dampening our ability to develop effective therapies to reduce their burden.

Here, we turn our attention towards heavy alcohol use and AUD, combining whole-brain structural, functional, and pharmacological information from diffusion MRI (dMRI), functional MRI (fMRI), and PET to investigate brain dynamics in individuals from the Human Connectome Project’s Young Adult dataset [13]. Using the brain’s structural (white matter) network as a guide, network control theory (NCT) [14] enables mapping of the brain’s dynamic state space by quantifying the energy required to transition between functional states. This type of energy can be referred to as *control* or *transition* energy. Recent work has utilized these tools to demonstrate that although the resting human brain has a spontaneous tendency to prefer certain brain state transitions over others, cognitive demands can overcome this tendency in a way that is associated with age and cognitive performance [15–17]. NCT has proven useful in describing brain dynamics in various cognitive states [15, 18], neuropsychiatric/degenerative conditions [16, 17, 19–21], and development [22, 23]. Importantly, NCT has also captured changes in brain dynamics due to neuromodulation [17, 24–26]. One such fMRI study showed increased transition energy under the D2 antagonist amulsipride compared to placebo [17]. They also showed that transition energy was negatively correlated with genetically predicted D2 receptor concentration, indicating those likely to have lower concentration of D2 receptors also had higher transition energy. This evidence supports the use of NCT to reveal shifts in the brain’s energetic landscape in response to receptor modulation/concentration, and, importantly, the hypothesis that decreased dopamine receptor function/concentration, as is known to occur in AUD, results in increased energetic demand to travel through the brain’s state space (i.e. increased transition energy). We thus propose using NCT as a unifying computational modeling approach that incorporates the effect of white matter and/or dopamine receptor differences in individuals with heavy alcohol use on their brain activity dynamics, with the goal of understanding neurobiological mechanisms of AUD at the whole brain level.

We utilize a network control framework to better our understanding of how brain structure and function is altered in heavy alcohol use. Using functional brain states from resting-state fMRI and the brain’s structural connectivity (SC) from dMRI, we compare transition energy in individuals with AUD and current heavy use of alcohol to that of individuals with minimal use of substances. We further relate these shifts in energetic demands to the complexity of brain activity, a well-known biomarker of information processing and brain health [27]. Then, to investigate changes in top-down and bottom-up signaling, we investigate how white matter differences in heavy alcohol use might alter signal propagation between subcortical structures and the frontoparietal network (FPN). Finally, we incorporate D2 receptor densities from PET to build a modeling framework that simulates dysfunction of the dopamine system and provides evidence for a mechanistic explanation for our observed findings.

## 2 Methods and Materials

### 2.1 Participants

We used data from (*n* = 958, 516 female, age = 28.73 (3.74 s.d.) years) participants of the Human Connectome Project - Young Adult S1200 [13] release. Individuals were assigned to the AUD group (*n* = 130, 30 female, age = 28 (3.7 s.d.) years) if they had a diagnosis of alcohol dependence/abuse, or were binge drinkers (*>*5 drinks per day at least weekly for the past year) and also reported having *>*3 drinks per day on average over the last year. Non-SUD individuals (*n* = 308, 213 female, age = 29 (3.8 s.d.) years) were individuals who did not have a diagnosis of any substance use disorder, and were not binge drinkers, and reported having *<*2 drinks per day on average for the past year. s.d. = standard deviation.

### 2.2 MRI data and preprocessing

We used publicly available, high resolution, preprocessed MRI data from the Human Connectome Project – Young Adult S1200 [13] in this study. HCP MRI data were acquired on a Siemens Skyra 3T scanner at Washington University in St. Louis. We examined resting-state functional MRI (2.0 mm isotropic, TR/TE = 720/33.1 ms, 8x multiband acceleration) from four 15 minute sessions and diffusion MRI (1.25 mm isotropic, TR/TE = 5520/89.5 ms, 3x multiband acceleration, b=1000, 2000, 3000 with 90 directions/shell)-both were collected with left-right and right-left phase encoding. Full preprocessing details were previously described in Gu et al.[28] in detail, and we summarize briefly here. Time series were denoised to remove signals from white-matter, CSF, and global gray-matter signal, and a high-pass filter removed signal *<*0.008Hz. The first ten frames of each scan were discarded to remove artifacts from scanner start up. For rsfMRI, outlier TRs identified based on head motion and global signal were replaced with linearly interpolated time-points. Preprocessed dMRI was further processed by multi-shell, multi-tissue constrained spherical deconvolution (CSD, [29]) and deterministic tractography (SD STREAM [30]) using MRtrix3, with SIFT2 global streamline weighting [31] and regional volume normalization. Regional time-series and structural connectomes for 958 HCP subjects were extracted using the 268-region Shen atlas [32].

### 2.3 Identification of brain-states

We concatenated the regional BOLD fMRI time-series from all 958 HCP-YA participants and performed *k*-means clustering with 20 repetitions and a maximum of 500 iterizations per repetition. Pearson’s correlation was used as the distance metric. *k* = 4 was chosen as the number of clusters based on prior work [24] and to allow calculation of MSC (Section 3.6). For each of the 438 individuals (130 AUD, 308 non-SUD) in the present analysis, brain-states were identified as the cluster centroids taken from all four of their fMRI scans.

### 2.4 Transition probability and state transitions

Using the partition of brain-states from *k* -means clustering, we calculated transition probabilities for each individual as the probability that any given state *i* was followed by state *j*. The number of state transitions for each individual was calculated as the number of times that any given state *i* was followed in the next volume by any state *j* where *j ̸*= *i*. These metrics were calculated separately for each fMRI scan and then averaged across scans prior to comparison.

### 2.5 Transition energy

We utilized each individual’s cluster centroids from Section 2.3 as brain-states to quantify state transition energies using NCT. Transition energy here is defined as the minimum energy input into a network—here, the structural connectome— required to move from one state to another [14, 33, 34]. To model neural dynamics, we used a linear time-invariant model:

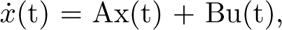

where A is an individual’s NxN structural connectivity matrix (normalized by its maximum eigenvalue plus 1 and subtracted by the identity matrix to create a continuous system) [34], x(t) is the regional activation at time t, B is the NxN matrix of control points, and u(t) is the external input into the system. Here, N is the number of regions in our parcellation. We selected *T* = 1 for the time-horizon, as in previous studies [15, 20, 22, 24, 25, 33]. Integrating u(t) over the time-horizon for a given transition yields the total amount of input that was injected into each region to complete the transition between states, and summing that value over all regions then gives the total amount of energy necessary to be injected over the whole brain. This summation represents *transition energy*. We calculated the pairwise transition energies between each of the four brain-states for each individual using this framework, using the identity matrix as the control strategy, B. In cases where there initial and target state were the same (Figure 4c, diagonal), transition energy was the energy required to maintain that state (i.e. resist the natural diffusion of activity through the SC). Average transition energy for each individual was calculated as the mean over all transitions. We also calculated transition energies between the subcortex and FPN using this framework, again using the identity matrix as the control strategy, B. For these calculations we constructed binary states where 1’s were assigned to brain regions belonging to the subcortex for the subcortical state, and the FPN [35] for the FPN state, and 0 elsewhere. For the transition energy calculations in the simulated dopamine dysfunction model, see Section 2.7.

### 2.6 Meta-state complexity

We calculated the meta-state complexity (MSC) of each individual’s *k* -means partition as previously described [24, 36]. In short, each individual’s partition was binarized based on assignment to either of the pairs of anticorrelated states (VIS-/+ or DMN-/+) to construct the meta-state time-series (Figure 3a). We then used the Lempel-Ziv algorithm (LZ76) [37] to quantify the compressability of, or information contained in, each binary meta-state time-series. This metric was calculated individually for each fMRI scan and then averaged across scans prior to comparison.

### 2.7 Simulated dopamine dysfunction

To simulate the impacts of decreased D2 receptor functioning on control energy, we began with all non-SUD individuals’ SCs and brain-states and recalculated average transition energies in a series of receptor-informed [24, 25, 38] scenarios by modifying the control strategy represented in the matrix B. First, to simulate the D2 receptor’s influence over average transition energy, we added rank-normalized D2 receptor densities (measured via PET-derived receptor binding potential) along the diagonal of B. The D2 receptor densities were obtained from a weighted average of 92 subjects from two D2 PET studies using the same tracer (FLB457) [39, 40] and compiled by Hansen et al [41]. We then compared these average transition energies against those obtained from increasing amounts of perturbation to the original receptor map. Specifically, we recalculated average transition energy using rank-normalized maps obtained after reducing the amount of D2 receptor density in the most D2-abundant regions (*>*95th percentile) by 20, 30, 40, and 50%. Each map was rank-normalized prior to addition to the B matrix in order to maintain the same overall amount of control given to the system and isolate only the effect of changes to the spatial allotment of control [33].

### 2.8 Statistical Comparisons

All between-group comparisons involving fMRI data (Figure 2b,d,e,f; Figure 3b) were made using ANOVAs controlling for age, sex, age:sex interaction, and fMRI in-scanner motion (average frame-wise displacement (FD)). Between group-comparisons investigating SC differences alone (Figure 4c,d) were made using the same ANOVA design as above sans fMRI in-scanner motion. Full tables for ANOVA results are located in the Supplementary Information. Correlations between average TE and the number of state-transitions (Figure 2g) and MSC (Figure 3c) were calculated using Spearman’s rank-correlation and p-values were obtained from permutation testing. The comparison between FPN to subcortex TE and subcortex to FPN TE (Figure 4b) was performed using both groups of participants and a paired t-test. Finally, the comparison of average TE obtained using the true D2 receptor map as a control strategy versus deplete maps were made using paired t-tests. All p-values were corrected for multiple comparisons using the Benjamini-Hochberg method where indicated (pFDR).

## 3 Results

### 3.1 Group Definitions

Non-SUD individuals were defined as persons without any substance use disorder, that were not binge drinkers (*>*5 drinks per day at least once weekly for the past year) and reported having *<*2 drinks per day on average for the past year. Subjects were collected into the AUD group if they had a diagnosis of alcohol dependence/abuse, or were binge drinkers, and also reported having *>*3 drinks per day on average over the past year. This last inclusion criteria allowed us to isolate the AUD group to those with current heavy use of alcohol. Between group comparisons were made using ANOVAs controlling for age, sex, age:sex interaction, and in-scanner motion (average frame-wise displacement) (see Section 2.8).

**Table 1:** Group classification table.

| Group | Females | Males | Total |
| --- | --- | --- | --- |
| AUD | 30 | 100 | 130 |
| nonSUD | 213 | 95 | 308 |

### 3.2 Commonly recurring patterns of brain activity

Data-driven clustering of all subjects’ regional BOLD fMRI time-series revealed four commonly recurring patterns of brain activity (Figure 1) that we operationalize as brain-states herein. The identified brain-states consisted of two pairs of anticorrelated activity patterns (i.e. meta-states), the first dominated by low and high-amplitude activity in the visual network (VIS-/+), and the second by low and high-amplitude activity in the default mode network (DMN-/+).

**Figure 1:**
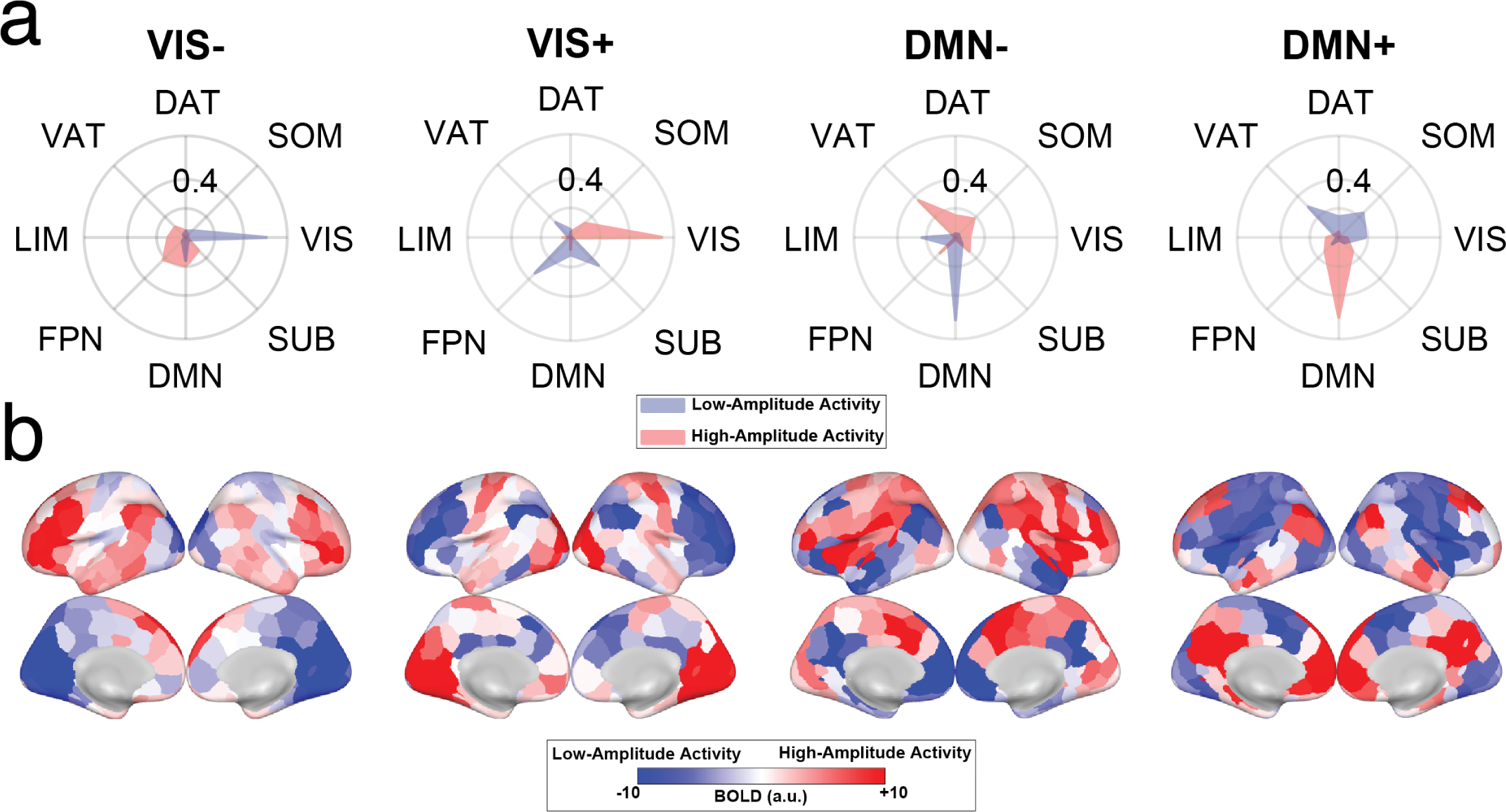
Four commonly recurring patterns of brain activity (brain-states) were identified using k-means clustering. Displayed are the group-average centroids. (a) Cosine similarity with canonical resting-state networks [35] was calculated for the positive (high-amplitude) and negative (low-amplitude) components separately for each brain-state. Each brain-state is labeled by its maximal cosine similarity value. (b) Mean BOLD activation of each brain-state plotted on the cortical surface. a.u. = arbitrary units. SUB - subcortical structures, VIS - visual network, SOM - somatomotor network, DAT - dorsal attention network, VAT - ventral attention network, LIM - limbic network, FPN - frontoparietal network, DMN - default mode network.

### 3.3 Less dynamic brain activity paired with larger transition energies in AUD

We calculated pairwise transition probabilities between each of the four brain-states (Figure 2a). Individuals with AUD showed a trend for lower likelihood of transitioning out of the DMN-state into the VIS- (*F* = 4.92, uncorrected p = 0.0389, pFDR = 0.210) and VIS+ (*F* = 4.27, uncorrected p = 0.0384, pFDR = 0.210) states, and a higher likelihood of staying in the DMN-state (*F* = 7.3, uncorrected p = 0.007, pFDR = 0.116) (Figure 2b), although none of these effects were significant after multiple comparisons correction. In general, individuals with AUD had fewer state transitions on average compared with non-SUD individuals (*F* = 7.24, pFDR = 0.0111) (Figure 2e). Applying network control theory to participants’ structural connectomes, we also calculated the minimum control energy, or *transition energy*, between each of the four brain-states for each individual (Figure 2c). Individuals with AUD showed higher transition energies for nearly every transition except for those into the DMN+ state (Figure 2d). Averaging across all pairwise transitions, AUD individuals also had larger average transition energy compared to non-SUD individuals (Figure 2f). Finally, across the entire group, individuals with larger average transition energy had fewer observed state transitions (rho = -0.77, pFDR *<* 0.0001) (Figure 2g).

**Figure 2:**
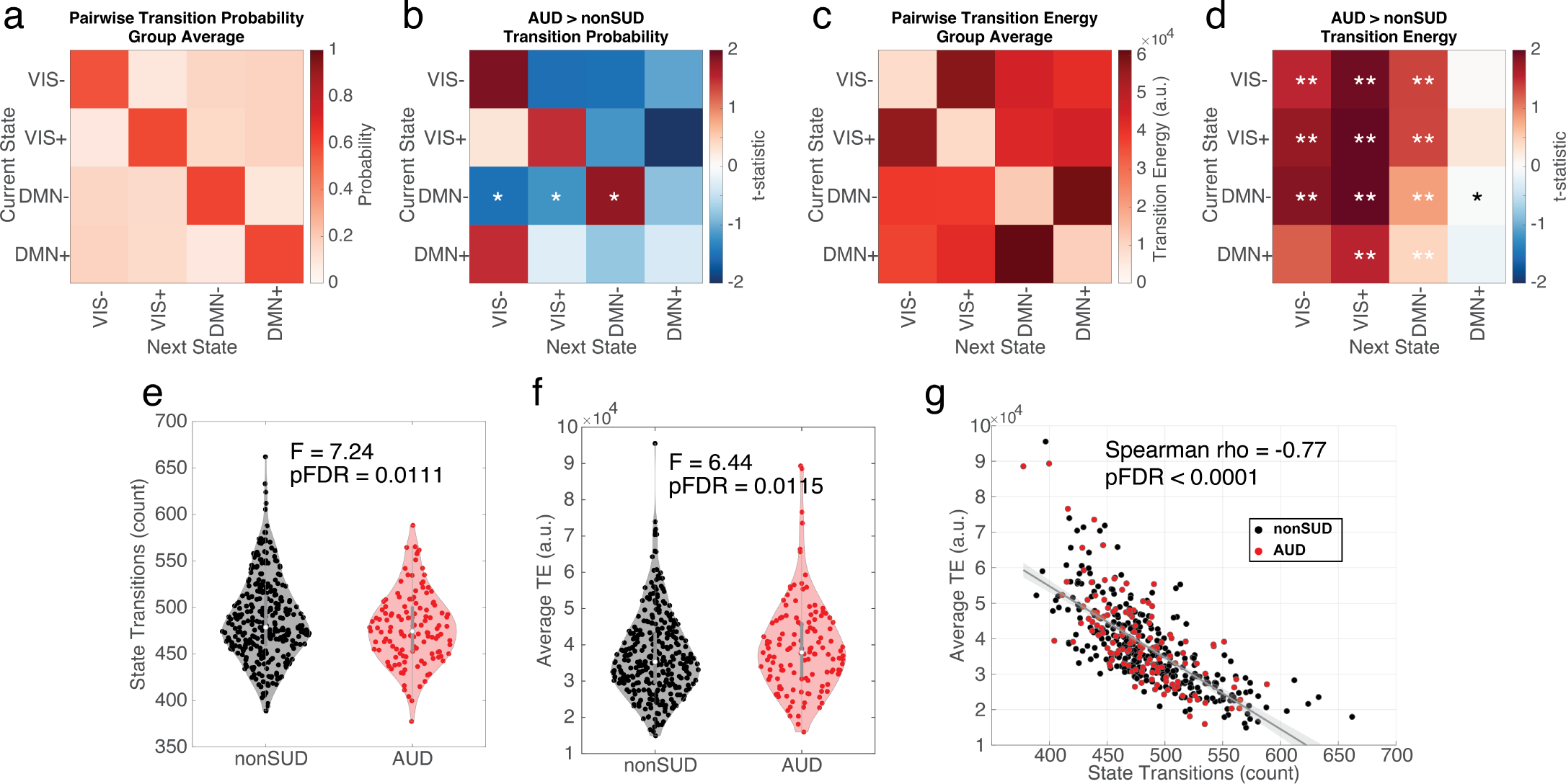
(a) Group-averaged pairwise transition probabilities observed between the four brain-states. (b) A trending group-effect for AUD on pairwise transition probabilities was observed for transitions out of DMN- and into VIS+/- and for maintaining the DMN-state. (c) Group-averaged pairwise transition energies. (d) Individuals with AUD had larger transition energies for the majority of potential state transitions. (e) There were overall fewer state transition observed in individuals with AUD. (f) The average transition energy across all transitions was larger in individuals with AUD. (g) Average transition energy was negatively correlated with the number of empirically observed state transitions on an individual level. In (b) and (d), t-statistics are visualized to illustrate the direction, however asterisks still represent p-values obtained from ANOVAs. * uncorrected p *<* 0.05; ** pFDR *<* 0.05. TE = transition energy. a.u. = arbitrary units.

While brain activity in AUD patients was less dynamic in terms of having fewer observed state-transitions, this is not a measure of brain activity complexity or information content. To this end, we next computed the meta-state complexity (MSC) of individuals’ brain-state time-series (Figure 3a). Individuals with AUD showed lower MSC compared to individuals without SUD (*F* = 10.92, pFDR = 0.0031) (Figure 3b), and average transition energy was negatively correlated with MSC across individuals (rho = -0.63, pFDR *<* 0.0001) (Figure 3c).

**Figure 3:**
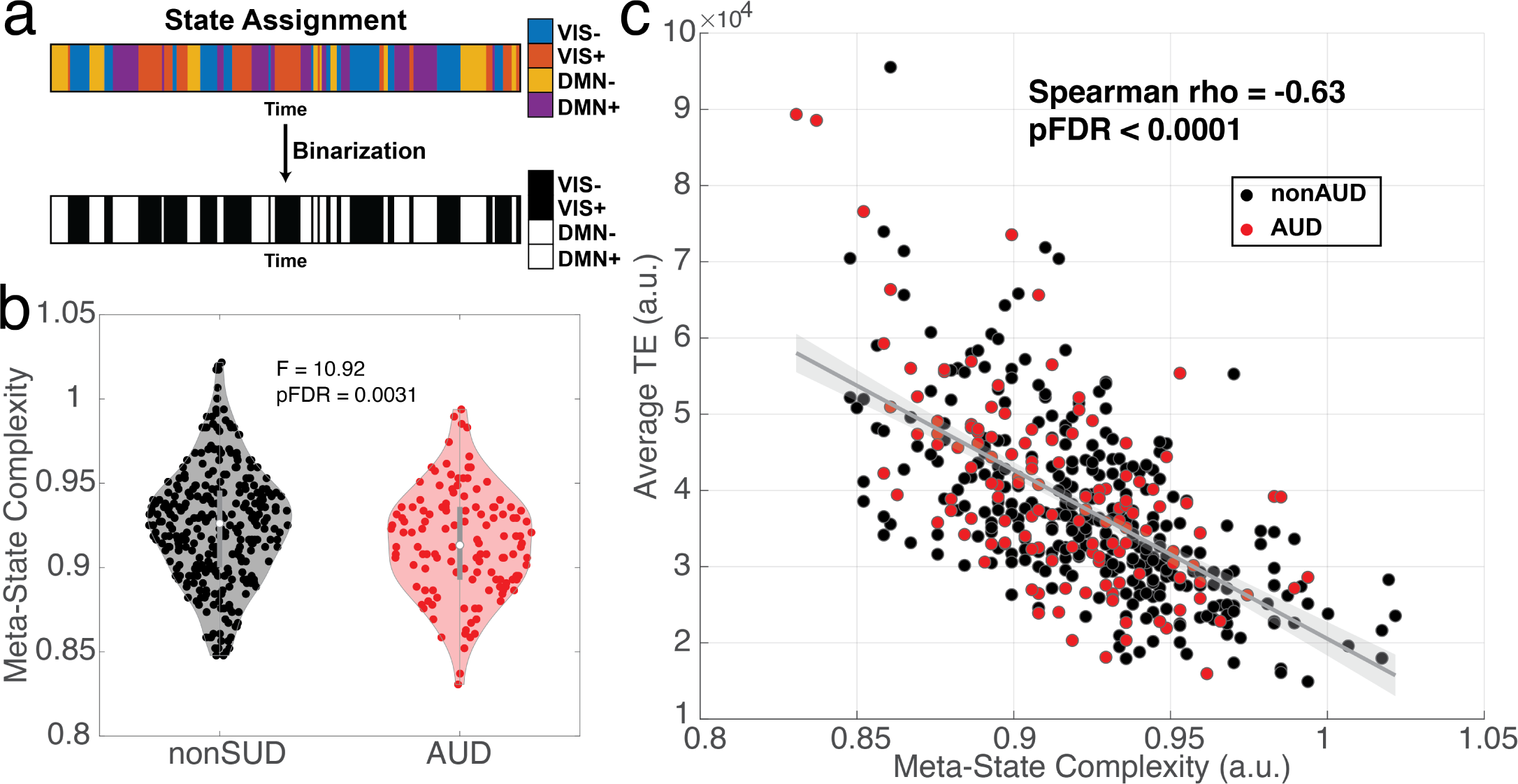
(a) Each participant’s partition of brain-states obtained from k-means clustering was binarized based on assignment to either VIS-dominated or DMN-dominated states. (b) Lempel-Ziv compressibility was run on the binarized sequences to characterize the complexity of the brain-state sequences (meta-state complexity; MSC). Individuals with AUD had significantly lower meta-state complexity compared to non-AUD individuals. (c) On an individual level, MSC and average transition energy were negatively correlated.

### 3.4 Higher subcortex to FPN transition energies in AUD

We next turned our attention towards transition energies between canonical subcortical and FPN states (Figure 4a) in order to test for asymmetrical communication patterns between these two parts of the brain in individuals with and without AUD. Due to homogeneous state definition across individuals (Section 2.5), this analysis is only revealing differences driven by changes in the white matter SC network. For all individuals, it required less energy to transition from the FPN to the subcortex than it did to transition in the reverse direction (*t* = -112, pFDR ¡ 0.0001) (Figure 4b). There was no group difference in transition energy between individuals with AUD and non-SUD individuals for the transition for the transition from the FPN to the subcortex (*F* = 1.63, pFDR = 0.2027) (Figure 4c). However, individuals with AUD did have larger TE for the transition from subcortex to FPN (*F* = 6.04, pFDR = 0.0216) (Figure 4d). Considering the direction of both trends, we performed a post-hoc evaluation of AUD’s effect on the TE asymmetry of these two transitions–that is–do individuals with AUD have a larger delta for transitioning one direction (FPN to subcortex) versus the other direction (subcortex to FPN)? Here, there was a slight trend suggesting that individuals with AUD have a larger TE asymmetry (*F* = 2.83, uncorrect p = 0.0934).

**Figure 4:**
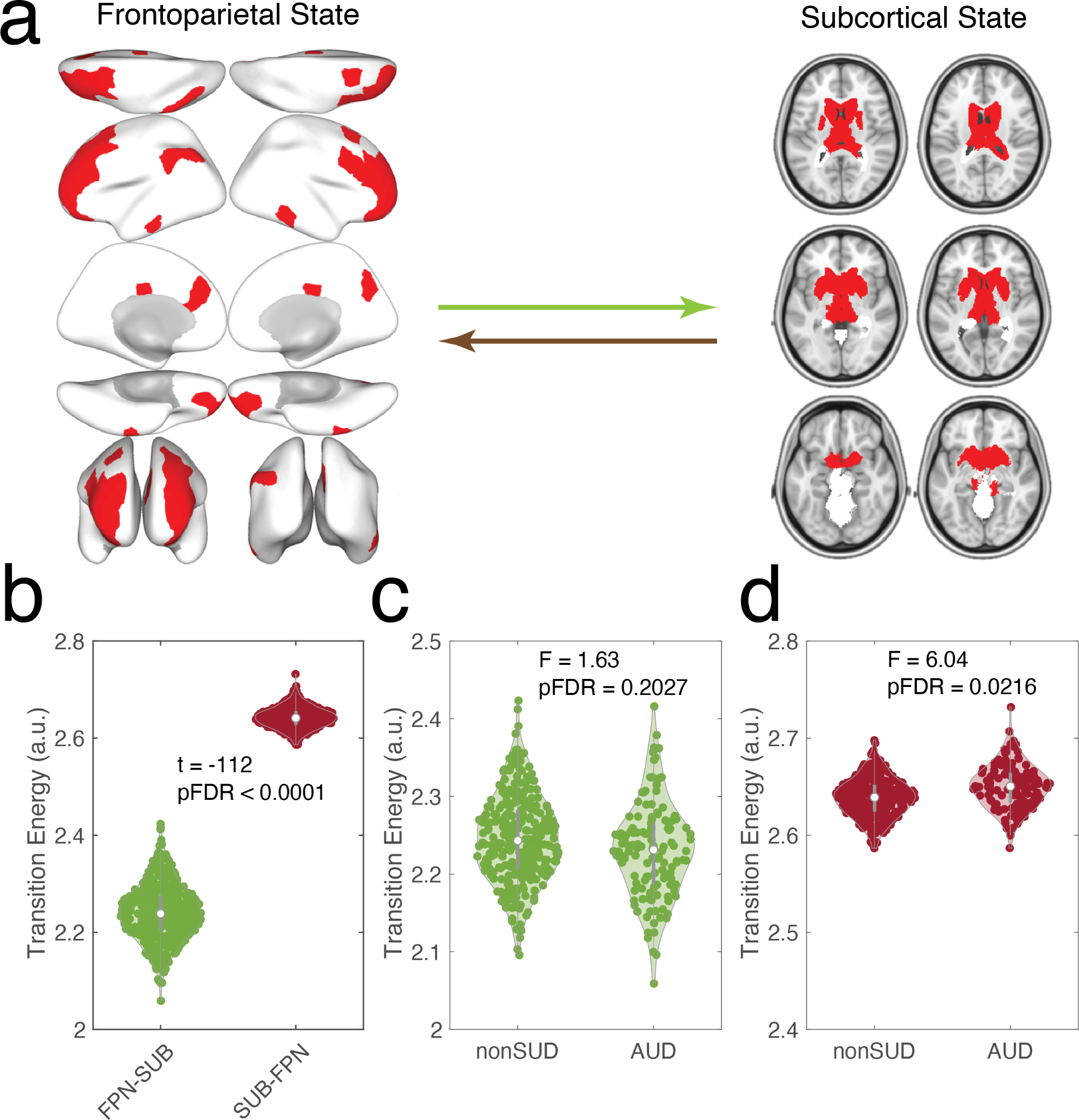
(a) Transition energies between canonical states of the frontoparietal network (FPN) and subcortical regions. (b) Across all subjects, it was more difficult to transition from the subcortex to the FPN (up the hierarchy) than it was to transition in the reverse direction (down the hierarchy). (c) There was no group effect on transitioning from the FPN to the subcortical network. (d) Individuals with AUD required more energy to transition from the subcortex to the FPN than those without an SUD.

### 3.5 Simulated dopamine dysfunction results in increased average transition energy

We deployed an *in silico* paradigm for studying the impacts of depleted dopamine receptor availability on transition energy (Figure 5). We simulated energies associated with *typical* dopaminergic functioning by calculating the average transition energy for non-SUD individuals using control weights derived from regional D2 receptor density maps (derived from PET scans in a separate population). We then assessed the impacts of D2 receptor depletion by recalculating average transition energy with a series of perturbed receptor maps and comparing the average transition energy from the perturbed D2 receptor maps to that of the true D2 receptor map (Figure 5a). We found that depleting the regions with the highest density of D2 receptors (*>*95th percentile which are mostly regions in the dorsal striatum), by 20 (*t* = 10.4, pFDR *<* 0.0001), 30 (*t* = 20.4, pFDR *<* 0.0001), 40 (*t* = 21.5, pFDR *<* 0.0001), and 50% (*t* = 8.5, pFDR *<* 0.0001) resulted in significant transition energy increases compared to the original, unperturbed map (Figure 5b).

**Figure 5:**
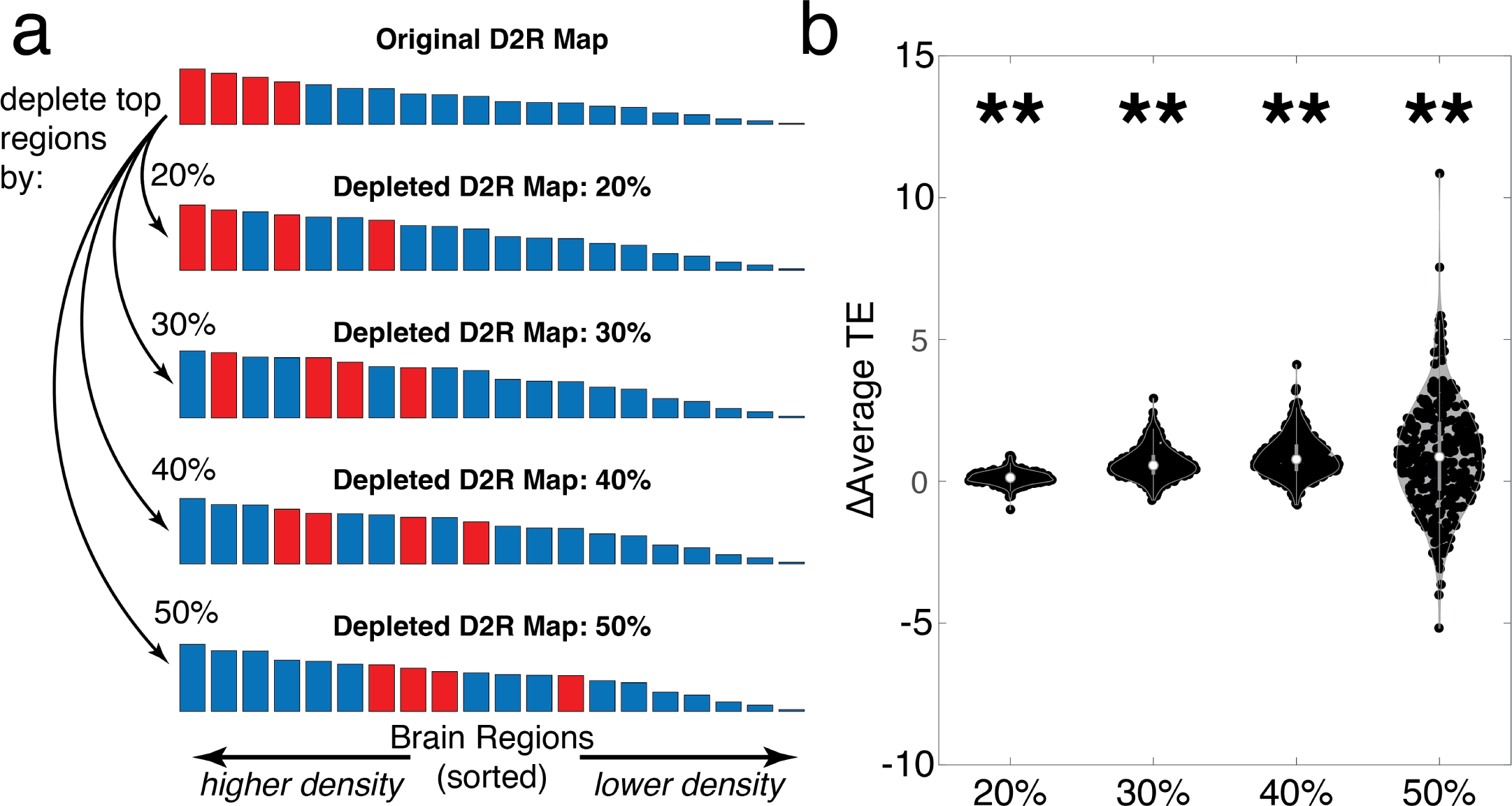
D2 receptor depletion simulation paradigm. (a) Top: the original PET-derived D2 receptor map ordered by the average density of D2 receptor availability per region (20 randomly selected regions shown for illustration purposes). To simulate dopamine receptor depletion or dysfunction, regions above the 95th percentile of D2 receptor density - mostly in the dorsal striatum, are depleted from their original values by 20, 30, 40, and 50%. Each of these maps were then used as control weights for calculating average TE for non-SUD individuals and the results of each depleted map was compared against those from the original map. (b) Each depleted map resulted in an increase in average transition energy compared to the original map. D2R = D2 receptor. ** pFDR *<* 0.0001

## 4 Discussion

We applied network control theory to understand how heavy current alcohol use alters both structure and function of the human brain in 438 individuals. Using individuals’ structural connectivity networks from dMRI and functional states from fMRI data, we found that transition energy in the brain was higher in individuals who have heavy current alcohol use (AUD) compared to those with minimal use of substances (non-SUD) (Figure 2d,f). Higher transition energy in AUD occurred alongside a concomitant decrease in the number of state transitions (Figure 2e) and MSC measured with resting-state fMRI (Figure 3b). Additionally, both the number of state transitions and MSC were strongly anti-correlated with average transition energy across all subjects (Figure 2g; Figure 3c). Using canonical states implicated in substance use, we found that AUD individuals required more energy to transition from the subcortex to the FPN (Figure 4d). Finally, we found that increasing the amount of dopamine dysfunction (by shifting control away from dorsal striatum regions with high D2 receptor expression), increased transition energies (Figure 5), mirroring the empirical results observed in AUD.

Network control theory is a computational framework that enables the quantification of state transition energies in the brain [14]. Transitions are modeled as a diffusion of initial states through the brain’s structural connectome, with energy being injected at each node (brain region) to control the trajectory toward the desired final state. The integration of these inputs over the length of the trajectory comprise the *control energy*, which we refer to simply as *transition energy*. Here, we calculated the transition energy between four commonly recurring patterns of brain activity in the resting-state fMRI time-series of each individual (Figure 1). Consistent with our hypothesis, individuals in the AUD group had larger transition energies compared to non-SUD individuals (Figure 2d,f). In addition, state-transitions and MSC were decreased in individuals with AUD compared to non-SUD. These findings suggest that brain dynamics under substance use, specifically alcohol, reflect a system entrenched in a state of low complexity and decreased information processing [27].

Brain entropy, here assessed via MSC, has been shown to index different states of consciousness as well as various brain disorders [24, 27, 36]. Brain entropy is impacted by the acute and/or chronic administration of various substances including alcohol [42], caffeine [43], nicotine [44], cocaine [45], as well as the psychedelics LSD, psilocybin, and DMT [36, 46–48]. Sevel et al [42] found that the acute administration of alcohol in healthy drinkers decreases brain entropy, a result that matches the sub-acute effects observed here in chronic heavy users of alcohol. Given that our energy and entropy results mirrored one another, we formally tested their association by correlating average transition energy and MSC across individuals (Figure 3c), and found a significant negative correlation. This relationship is consistent with previous studies showing an inverse relationship between transition energy and entropy that is modulated by disease [20] and pharmacological intervention [24, 25].

Previous work suggests that network control theory can capture structural differences relevant for executive functioning and development [22, 49]. Cui et al [49] demonstrated that the amount of energy required to activate the FPN decreases throughout development, and additionally, that individuals who required less energy to activate the FPN had higher executive functioning. Here, we studied the bi-directional transitions between the subcortex and the FPN (Figure 4) due to the known involvement of dopaminergic mesocorticaolimbic signaling pathways and fronto-subcortical circuits in addiction [10, 50–52]. We found that AUD individuals required more energy to transition from the subcortex to the FPN than non-SUD individuals (Figure 4d). This finding suggests that the coarse-grained structural connectome topology of individuals with AUD is organized in a way that limits the natural diffusion of information from subcortical structures to the FPN. This possibly relates to atrophy of regions belonging to corticostriatal-limbic circuits observed in AUD [53, 54] or increased difficulty in activating the FPN which could be associated with decreased executive functioning found in AUD [55].

Individuals with AUD show reduced levels of D2 receptors in subcortical limbic and striatal areas, which is also where D2 receptors are most dominantly expressed [4, 56, 57]. We developed an *in silico* D2 receptor depletion model in order to test the correspondence between spatial patterns of aberrant dopaminergic signaling and our observation of increased transition energies in the AUD group. We recalculated average transition energy in non-SUD individuals using five different sets of control weights corresponding to increasing amounts of disruption to typical D2 receptor signaling. We found that reducing the amount of control given to the regions most richly expressed in D2 receptors increased transition energies (Figure 5). This suggests a potential link between decreased D2 receptor functioning and larger transition energy in AUD. Indeed, prior work has demonstrated increased transition energies in individuals administered a D2 antagonist and negative correlations between genetically estimated D2 receptor densities and global transition energies [17].

Due to the limitations of the available data, we were not able to take into account important factors for substance use such as the amount of time since the most recent drink or the duration of alcohol use. It is possible that heterogeneity in influential factors such as the severity of dependence and active states of substance use (withdrawal or intoxication) could impact an individual’s cognitive state and thus the amount of vigilance/cortical arousal during fMRI scanning, which is known to influence properties such as signal amplitude [58]. While we controlled for sex, age, and sex:age interactions in our results, we did not seek to formally evaluate these relationships here; we will do so in future work.

We combined dMRI, fMRI, and PET to perform a whole-brain evaluation of alcohol use disorder’s impacts on human brain structure and function. We found functional landscapes in AUD were reflective of less dynamic and complex activity, with greater barriers to transition between brain-states compared to individuals without an SUD. We also found higher energetic demands to propagate signals through the structural connectome from the subcortex to the FPN in AUD, and, finally, evidence that dopamine receptor dysfunction could be a contributing mechanism to this increased energetic demand for state transitions in AUD. This study demonstrates the ability of this multi-modal NCT framework for uncovering shifts in brain dynamics and potentially for uncovering neurobiological mechanisms of these shifts. The latter understanding is key if we are to better diagnose, prevent, track and treat AUD so we can help reduce the individual and societal burden of this debilitating disorder.

## Acknowledgements

LS was supported by the National Institute on Drug Abuse of the National Institutes of Health under Award Number T32 DA03980. AIL acknowledges the support of the Natural Sciences and Engineering Research Council of Canada (NSERC), [funding reference number 202209BPF-489453-401636, Banting Postdoctoral Fellowship] and FRQNT Strategic Clusters Program (2020-RS4-265502 - Centre UNIQUE - Union Neuroscience & Artificial Intelligence - Quebec) via the UNIQUE Neuro-AI Excellence Award. LP was supported by the National Institute Of Mental Health of the National Institutes of Health under Award Number R00MH127296. AK was supported by the National Institute of Mental Health of the National Institutes of Health under Award Number RF1 MH123232.

