## Supplemental Information for "Altered structural connectivity and functional brain dynamics in individuals with heavy alcohol use"

### 1 Supplementary Tables

| Source | Sum Sq. | d.f. | Mean Sq. | F | Prob>F |
|---|---|---|---|---|---|
| AUD | 11376.1 | 1 | 11376.1 | 7.24 | 0.0074 |
| Sex | 199.2 | 1 | 199.2 | 0.13 | 0.722 |
| Age | 43373.5 | 1 | 43373.5 | 27.6 | 0 |
| Sex:Age | 49.5 | 1 | 49.5 | 0.03 | 0.8592 |
| mean framewise displacement | 53371.7 | 1 | 53371.7 | 33.96 | 0 |
| Error | 678927.4 | 432 | 1571.6 | | |
| Total | 805502.7 | 437 | | | |

Table 1: State-transitions (count) ANOVA

| Source | Sum Sq. | d.f. | Mean Sq. | F | Prob>F |
|---|---|---|---|---|---|
| AUD | 8.30e+08 | 1 | 8.30e+08 | 6.44 | 0.0115 |
| Sex | 6.08e+07 | 1 | 6.08e+07 | 0.47 | 0.4925 |
| Age | 5.27e+09 | 1 | 5.27e+09 | 40.91 | 0 |
| Sex:Age | 1.65e+08 | 1 | 1.65e+08 | 1.28 | 0.2584 |
| mean framewise displacement | 5.33e+07 | 1 | 5.33e+07 | 0.41 | 0.5205 |
| Error | 5.57e+10 | 432 | 1.29e+08 | | |
| Total | 6.23e+10 | 437 | | | |

Table 2: Average transition energy ANOVA

| Source | Sum Sq. | d.f. | Mean Sq. | F | Prob>F |
|---|---|---|---|---|---|
| AUD | 0.01073 | 1 | 0.01073 | 10.92 | 0.001 |
| Sex | 0.00003 | 1 | 0.00003 | 0.03 | 0.8598 |
| Age | 0.01782 | 1 | 0.01782 | 18.13 | 0 |
| Sex:Age | 0.00003 | 1 | 0.00003 | 0.03 | 0.8647 |
| mean framewise displacement | 0.03260 | 1 | 0.03260 | 33.16 | 0 |
| Error | 0.42464 | 432 | 0.00098 | | |
| Total | 0.49926 | 437 | | | |

Table 3: Meta-state complexity ANOVA

| Source | Sum Sq. | d.f. | Mean Sq. | F | Prob>F |
|---|---|---|---|---|---|
| AUD | 0.00521 | 1 | 0.00521 | 1.63 | 0.2027 |
| Sex | 0.00087 | 1 | 0.00087 | 0.27 | 0.6027 |
| Age | 0.04326 | 1 | 0.04326 | 13.51 | 0.0003 |
| Sex:Age | 0.00001 | 1 | 0.00001 | 0 | 0.9555 |
| Error | 1.38644 | 433 | 0.0032 | | |
| Total | 1.47023 | 437 | | | |

Table 4: FPN to SUB transition energy ANOVA

| Source | Sum Sq. | d.f. | Mean Sq. | F | Prob>F |
|---|---|---|---|---|---|
| AUD | 0.00235 | 1 | 0.00235 | 6.04 | 0.0144 |
| Sex | 0.00083 | 1 | 0.00083 | 2.13 | 0.1451 |
| Age | 0.0018 | 1 | 0.0018 | 4.64 | 0.0318 |
| Sex:Age | 0.00008 | 1 | 0.00008 | 0.21 | 0.6436 |
| Error | 0.1684 | 433 | 0.00039 | | |
| Total | 0.20038 | 437 | | | |

Table 5: SUB to FPN energy ANOVA

| Source | Sum Sq. | d.f. | Mean Sq. | F | Prob>F |
|---|---|---|---|---|---|
| AUD | 0.01456 | 1 | 0.01456 | 2.83 | 0.0934 |
| Sex | 0.00339 | 1 | 0.00339 | 0.66 | 0.4172 |
| Age | 0.06274 | 1 | 0.06274 | 12.18 | 0.0005 |
| Sex:Age | 0.00015 | 1 | 0.00015 | 0.03 | 0.8641 |
| Error | 2.22939 | 433 | 0.00515 | | |
| Total | 2.43244 | 437 | | | |

Table 6: TE Asymmetry (SUB to FPN - FPN to SUB) ANOVA
